# On the Structure and Mechanism of Two-Pore Channels

**DOI:** 10.1101/181578

**Authors:** Alexander F. Kintzer, Robert M. Stroud

**Affiliations:** University of California, San Francisco Department of Biochemistry and Biophysics

**Author notes:** Correspondence to Robert Stroud.

**Keywords:** Ion channels, Structure, Membrane protein, Lysosome, Transport

## Abstract

In eukaryotes, two-pore channels (TPC1-3) comprise a family of ion channels that regulate the conductance of Na^+^ and Ca^2+^ ions across cellular membranes. TPC1-3 form endolysosomal channels, but TPC3 can also function in the plasma membrane. TPC1/3 are voltage-gated channels, but TPC2 opens in response to binding endolysosome-specific lipid phosphatidylinositol-3,5-diphosphate (PI(3,5)P_2_). Filoviruses, such as Ebola, exploit TPC-mediated ion release as a means of escape from the endolysosome during infection. Antagonists that block TPC1/2 channel conductance abrogate filoviral infections. TPC1/2 form complexes with the mechanistic target of rapamycin complex 1 (mTORC1) at the endolysosomal surface that couple cellular metabolic state and cytosolic nutrient concentrations to the control of membrane potential and pH. We determined the X-ray structure of TPC1 from *Arabidopsis thaliana* (AtTPC1) to 2.87Å resolution–one of the two first reports of a TPC channel structure. Here we summarize these findings and the implications that the structure may have for understanding endolysosomal control mechanisms and their role in human health.

**Abbreviations:**

mTORC1: Mechanistic target of rapamycin complex 1
TPC: Two-pore channel
PI(3,5)P_2_: Phosphatidylinositol-3,5-diphosphate
AtTPC1: *Arabidopsis thaliana* TPC1
NED19: Trans-Ned-19
VSD: Voltage-sensing domain
P1: Pore domain in S5-S6
P2: Pore domain in S11-S12
Ca_v_: Voltage-gated calcium channel
Na_v_: Voltage-gated sodium channel
K_v_: Voltage-gated potassium channel
NTD: N-terminal domain
CTD: C-terminal domain
EF: EF-hand domain
NAADP: Nicotinic acid adenine dinucleotide phosphate
PI(4,5)P_2_: Phosphatidylinositol-4,5-diphosphate
DHP: Dihydropyridine
PAA: Phenylalkylamine
BTZ: Benzothiazepine
Ca_a_^2+^: Activating Ca^2+^-ion
Ca_i_^2+^: Inhibitory Ca^2+^-ion
fou2: Fatty acid oxygenation up-regulated 2
SLC38a9: Sodium-coupled neutral amino acid transporter 9
NPC1: Niemann-Pick C1
PKA: Protein kinase A
PKC: Protein kinase C
PKG: Protein kinase G
H^+^: ATPase - Proton Pump
^32^P: – Phosphorus-32

## Main Text

Endolysosomes facilitate cellular trafficking, nutrient uptake and release, protein degradation, and pathogen evasion[1]. These vital cellular functions hinge on the precise control of ionic (Na^+^, K^+^, Ca^2+^, Cl^-^, H^+^) gradients that define the endolysosomal membrane potential[2]. Ca^2+^-release and uptake through ion channels and transporters enables communication[3–5] between organelles and control of vesicular movement[6]. Transporters, proton pumps, and ion channels maintain the lysosomal pH required for proteolytic functions[7]. Despite the prevalence of human disorders that exhibit lysosome dysfunction[1,8], the molecular mechanisms that control lysosome transport remain poorly understood as compared with plasma membrane transport[9].

Two-pore channels (TPCs) form a subfamily (TPC1-3) of eukaryotic voltage- and ligand-gated cation channels[10,11]. TPC1/2 function as part of the ion transport machinery that regulate the endolysosomal resting membrane potential and pH[12]. TPC1/2 knockout mice have lower cellular amino acid levels and endurance during fasting[13], suggesting that TPC-mediated control of the endolysosome membrane potential and pH regulates nutrient transport. Complexes between TPC1/2 and the mechanistic target of rapamycin complex 1 (mTORC1) on the endolysosome link cellular metabolic status and nutrient levels to regulation of amino acid efflux[13]. MTORC1 inhibits channel opening, presumably by phosphorylation, though no sites have been identified. TPC1/2-mTORC1 complexes may be therapeutic targets because greater than thirty different mTOR mutations exist in human cancers, a subset of which activate mTORC1 and facilitate tumor proliferation and chemotherapy resistance[14]. Furthermore, mutations in TPC2 disrupt Na^+^/Ca^2+^ gradients in endolysosome-like melanocytes[15,16], which may contribute to the development of skin cancer[17]. TPCs may also be important drivers of lung cancer metastasis[18]. The molecular nature of TPC1/2-mTORC1 interactions demands further investigation. Filoviruses exploit TPC1/2 ion conductance to escape the endolysosome prior to degradation in the lysosome[19]. Pharmacological block of TPC1/2 by trans-Ned-19 (NED19) and certain calcium channel blockers inhibits viral infectivity. Antagonists could prevent viral uptake through effects on membrane fusion, receptor binding, or proteolytic activation.

To approach an atomic understanding of how TPC channels control endolysosomal ion transport, we determined the structure of TPC1 from *Arabidopsis thaliana* (AtTPC1) by X-ray crystallography to 2.87Å resolution (Figure 1, ref. [20]). A second report determined a 3.3Å structure of AtTPC1 along with electrophysiological measurements[21]. These were the first structures of a TPC channel. A single TPC1 polypeptide contains two tandem Shaker-like domains (D1 and D2, ref. [22,23]) that dimerize to form a central quasi-tetrameric channel. Each chain contains two voltage-sensing domains (VSDs) involving transmembrane segments S1-S4 (VSD1) and S7-S10 (VSD2), two pore domains in S5-S6 (P1) and S11-S12 (P2), and activation gates following S6 and S12. D1 and D2 share 20-30% sequence identity with particular domains in voltage-gated calcium (Ca_v_) and sodium channels (Na_v_), suggesting that TPCs are a family intermediate between the tetrameric channels, like voltage-gated potassium (K_v_) channels, and Ca_v_s/Na_v_s that have four pore-forming domains in series on a single chain[24]. This heritage might explain why some inhibitors of Ca_v_ channels also block TPC channels[19,25].

**Figure 1.**
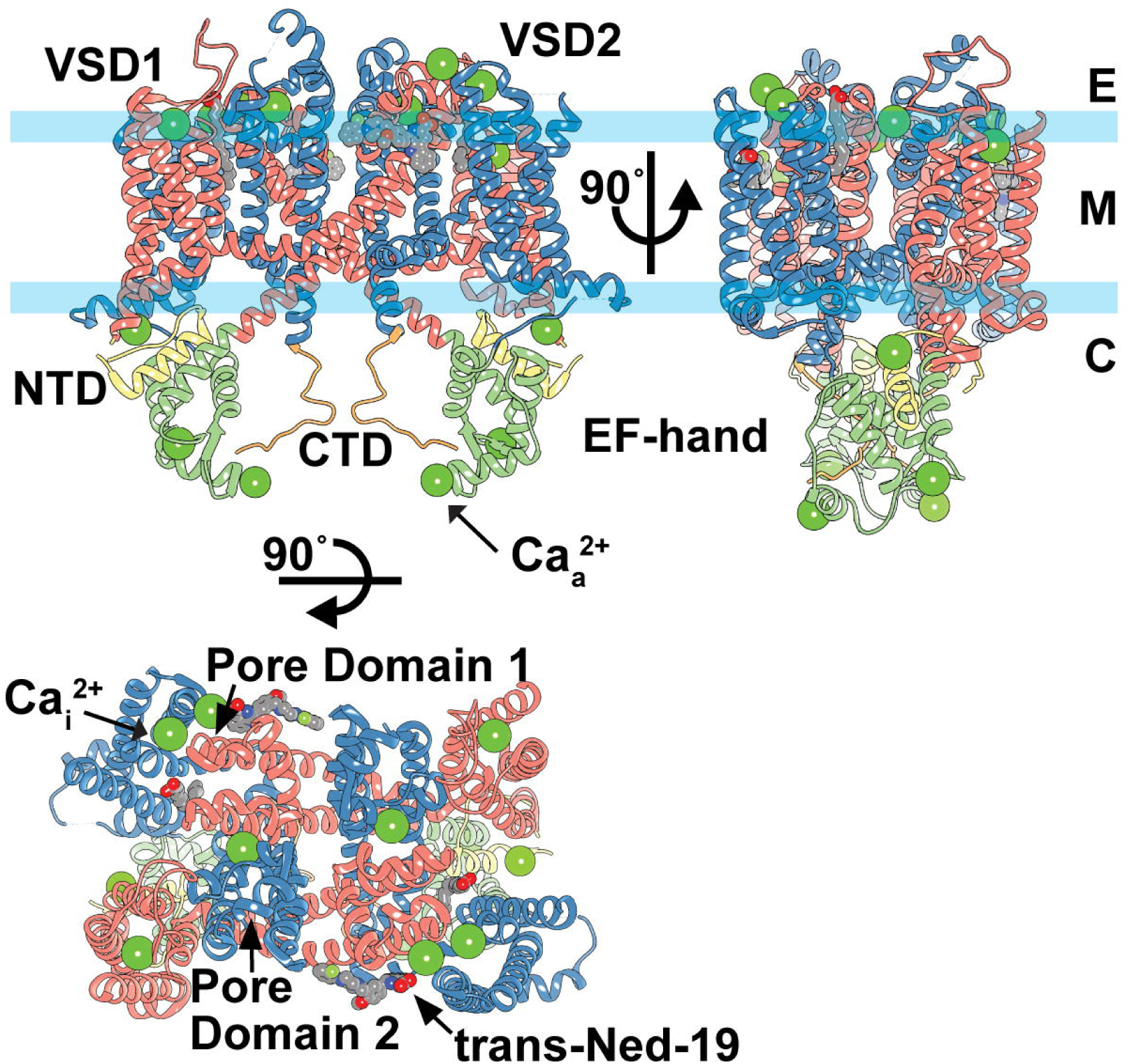
Overview of the AtTPC1 Structure. Views down (top) long and short channel axes, and (bottom) top-down through the central channel of AtTPC1 (PDB **5DQQ**; ref. [20]). Boundaries for endolysosome/vacuole (E), membrane (M), and cytoplasm (C) are shown. Ca^2+^-ions, including the sites for luminal inhibition (**Ca_i_^2+^**) and cytoplasmic activation (**Ca_a_ ^2+^**), are shown as green spheres.

The structure of AtTPC1 is asymmetric. Non-equivalence of D1 and D2 and tandem architecture create asymmetry in the molecule that stems from differing lengths of the lateral helices joining VSD1 to P1 (S4-S5) and VSD2 to P2 (S10-S11). S4 partially unravels at its base, causing S4-S5 to be shorter than S10-S11 by one helical turn. The angle between S4 and S5 (87°) is more obtuse than between S10 and S11 (38°), leading to elongation along the dimer axis and contraction along D1-D2 that result in a rectangular 2-fold symmetric channel. Four pore domains form the central channel in a 2-fold symmetric arrangement (P1-P2-P1-P2). Discussion in sections below highlight the potential roles of TPC channel asymmetry in voltage-dependent activation and ion selectivity.

Plant TPC1 channels contain cytosolic N-terminal domains (NTD), C-terminal domains (CTD), and two EF-hand domains (EF) between the two pore subunits that confer activation by Ca^2+^-ions[10,26]. A di-leucine motif in the NTD directs trafficking to the endolysosome (or vacuole in plants)[27]. Both the NTDs and CTDs have roles in channel activation; when they are removed, channels no longer function[28,29].

In most eukaryotic cells, but not in plants or yeast, nicotinic acid adenine dinucleotide phosphate (NAADP) triggers the release of Ca^2+^-ions from acidic intracellular stores, causing temporary increases in the cytosolic Ca^2+^-ion concentration[11,30], influencing biological processes ranging from cell differentiation to cardiac function[31–33]. Fluorescence Ca^2+^-imaging experiments suggest that NAADP triggers Ca^2+^-release from acidic Ca^2+^-stores containing TPC1-3[11,34–36]. However, photoaffinity labeling experiments with ^32^P-labeled NAADP show that it does not directly bind TPCs[37,38]. Rather, an unidentified NAADP-binding protein may be required for activation of TPCs[39]. It is also possible that other channels and/or other organelles may be the target and functional source of Ca^2+^-ions. Identification of the putative NAADP-receptor will be important for determining the role of TPC channels in endolysosomal Ca^2+^-signaling.

## Activation Mechanisms

TPC1 confers endolysosomes with electrical excitability[40]. TPC1 is primarily a voltage-gated channel, but luminal pH[40], cytosolic and luminal Ca^2+^-ions[41–43], and endolysosome-specific lipid PI(3,5)P_2_[12] regulate channel conductance. Mammalian TPC2 is voltage-insensitive and activated by PI(3,5)P_2_ and potentially by NAADP[11]. TPC3 is not present in humans, but the mammalian and vertebrate orthologues localize to both plasma membranes and endolysosomes[11,36,44]. On plasma membranes, fish and frog TPC3 behaves as voltage-gated, Na^+^-selective channels, insensitive to PI(3,5)P_2_ and PI(4,5)P_2_[44]. On endolysosomes, chicken and rat TPC3 may conduct Ca^2+^-ions in response to NAADP[36], whereas conflicting reports of invertebrate TPC3 NAADP-sensitivity exist[45,46].

Figure 2 summarizes the known or suggested activation/regulation mechanisms of TPCs[10,12,47–51]. Cytoplasmic Ca^2+^-ions are required for voltage-dependent activation of plant TPC1[10]. Gating of plant TPCs is likely a multistep process that requires coupling of the VSDs to the cytoplasmic EF domains. Each EF domain contains two Ca^2+^-binding sites (EF1 and EF2). Substitution of Ca^2+^-coordinating residues has shown that only EF2 is responsible for Ca^2+^-activation[26].

**Figure 2.**
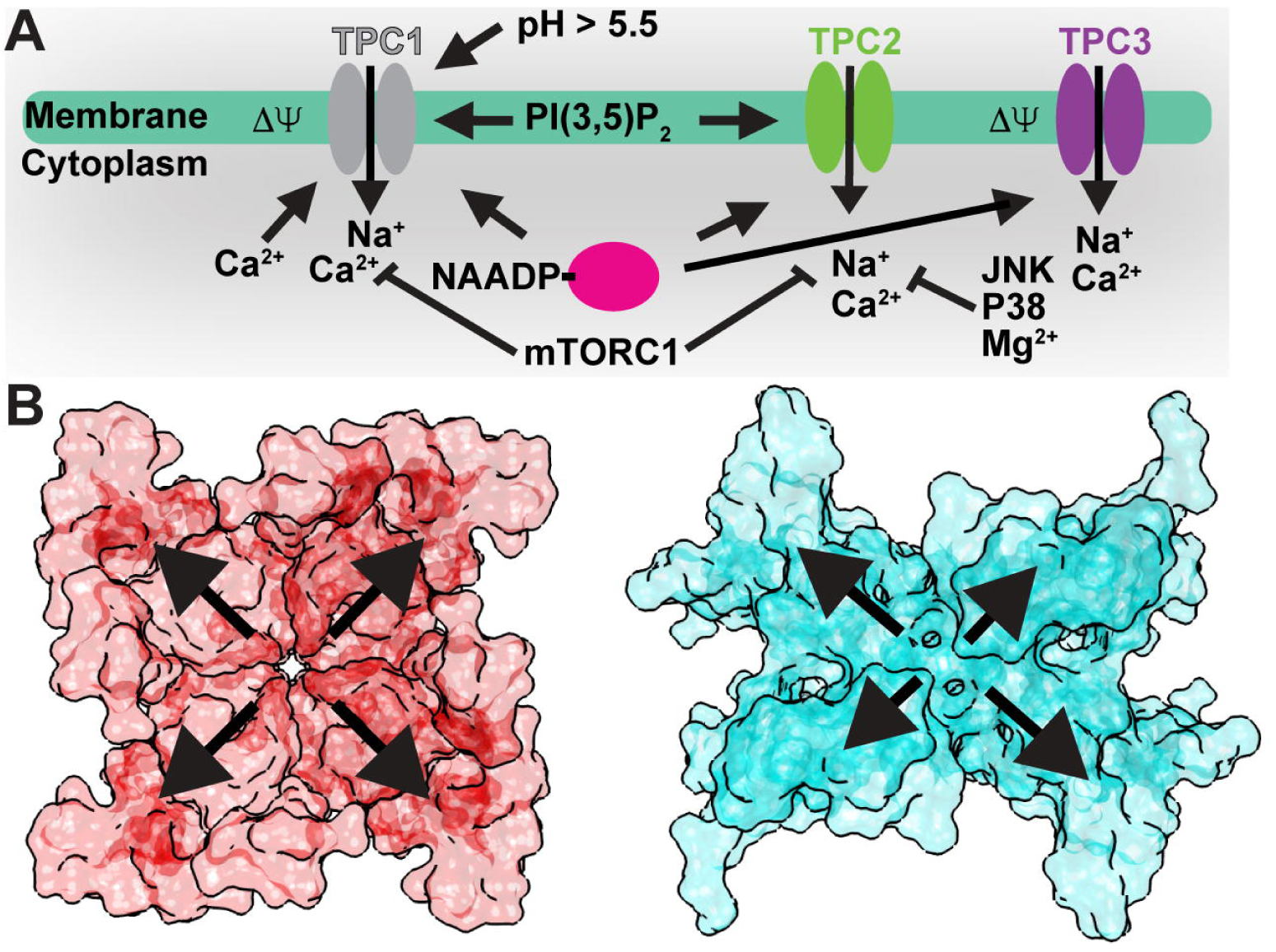
Figure 2. TPC Activation Mechanisms. A) Summary of luminal/extracellular and cytoplasmic agents and processes that modulate TPC channel opening. B) Surface renderings of a (left) symmetrical tetrameric channel (Na_v_Ab; PDB **3RVY**; ref. [56]) versus (right) the asymmetric tandem TPC1 dimer (AtTPC1; PDB **5DQQ**; ref. [20]). Outward arrows highlight potential differential contributions from the sensory and pore domains to channel activation.

How do the VSDs and EFs couple structurally to facilitate activation by both membrane potential and cytoplasmic Ca^2+^-ions? The AtTPC1 structure captures the Ca^2+^-activated state of the EF2[20]. Ca^2+^-bound EF2 forms an intramolecular complex with the CTD via a salt-bridge between D376 of EF2 and R700 of the CTD. CTD residue S701 coordinates the activating Ca^2+^-ion (Ca_a_ ^2+^; Figure 1) along with E374 and D377. The EF2-CTD interaction may serve as the ‘ Ca^2+^-switch’, coupling Ca^2+^-binding in EF2 and the CTD to activation. R552 in the S10-S11 linker of VSD2 forms a salt-bridge with D685 of the CTD that could link the conformation of EF2-CTD to voltage-sensing. Removal of cytosolic Ca^2+^ could weaken the EF2-CTD interaction, increase dynamics in the CTD, and disrupt conformational coupling to VSD2.

The EF2-CTD interactions are probably conserved between TPCs since the NTDs and CTDs facilitate channel activation in both mammalian and plant TPCs[28,29]. In human TPC2, the CTD mutation G734E correlates with changes in skin and hair pigmentation and the defective pigment synthesis observed in melanomas[17,52]. This suggests that the EF2-CTD interaction could be important for channel function in humans.

Luminal Ca^2+^-ions inhibit plant TPC1 channel activation at millimolar concentrations via binding sites in the S7-S8 loop of VSD2[47]. The AtTPC1 mutation D454N, named *fou2* (fatty acid oxygenation up-regulated 2), causes an increase in stress hormone jasmonate synthesis as part of the Ca^2+^-dependent wounding response in plants[53,54]. D454N increases channel open probability by shifting the voltage-dependence of VSD2 towards more depolarizing potentials and abolishes inhibition by luminal Ca^2+^-ions[47,21]. The structure of AtTPC1 and electrophysiological measurements clarified the mechanism of luminal Ca^2+^-inhibition by revealing two Ca^2+^-binding sites (site 1 and site 2) at the luminal face of VSD2[20,21]. The approximate location of these two Ca^2+^-sites had been predicted previously[55], though the models did not explain how either site could directly modulate voltage-sensing. The structure shows that D240 in P1, D454 in the S7-S8 loop, and E528 in S10 coordinate site 1, whereas E239 in P1 and E457 in the S7-S8 loop coordinate site 2. Site 1, not site 2, is the luminal inhibitory Ca^2+^ site because mutations in site 1 (D240N, D454N, E528Q), but not site 2 (D239N, E457Q), eliminate luminal Ca^2+^ inhibition (Ca_i_^2+^; Figure 1) in electrophysiological recordings[21,47]. Site 1 mutations also shift voltage-dependent activation by nearly +50mV, such that the channel is in an open-activated state under crystallization conditions (0mV)[21]. The function of Ca_i_^2+^ may be to tune the endolysosomal/vacuolar membrane potential by shifting voltage-dependent activation of TPC1. Fortuitously, this unique luminal Ca^2+^-site may also enable trapping of multiple conformational states of the channel by changing the Ca^2+^-concentration.

Wild-type AtTPC1 adopts a closed-resting state at 0mV in the presence of luminal millimolar concentrations of Ca^2+^-ions[21]. The functional voltage sensor VSD2 is the first case of the resting-state conformation with respect to other voltage-gated ion channel structures. The three functional gating charges (R1; R537, R2; R540, R3; R543) in S10 shift downward towards the cytosolic leaflet with respect to active-state structures (Figure 3) [56–59]. Comparing AtTPC1 resting-state and other active-state VSDs we see a striking change in the overall VSD structure. In the resting-state, the last two gating charges (R2, R3) of AtTPC1 are exposed to the cytoplasm, and the first gating charge R1 is at the well-conserved hydrophobic charge transfer center[60]. We deduce that during activation the movement of S4/S10 in VSD2 exposes the first two gating charges (R1, R2) to the endolysosomal lumen or extracellular space. This would electrically move 2 gating charges to the other side of the membrane per activation cycle per VSD2. Activation of both VSD2s would account for the estimated gating charge of ~3.9 for AtTPC1 to achieve unitary open probability[21]. Electrophysiological measurements also confirm that VSD2 is the active voltage-sensor and VSD1 is voltage-insensitive[21,61]. Movements in S7 and S9 lead to an overall closing of the solvent vestibule to the gating charges at the cytoplasmic face of VSD2, accommodating the movement of S10 during activation and opening the solvent to gating charges via the endolysosomal/extracellular face. This effectively moves gating charge across the membrane barrier and mechanically displaces the S10-S11 linker to open the channel activation gates.

**Figure 3.**
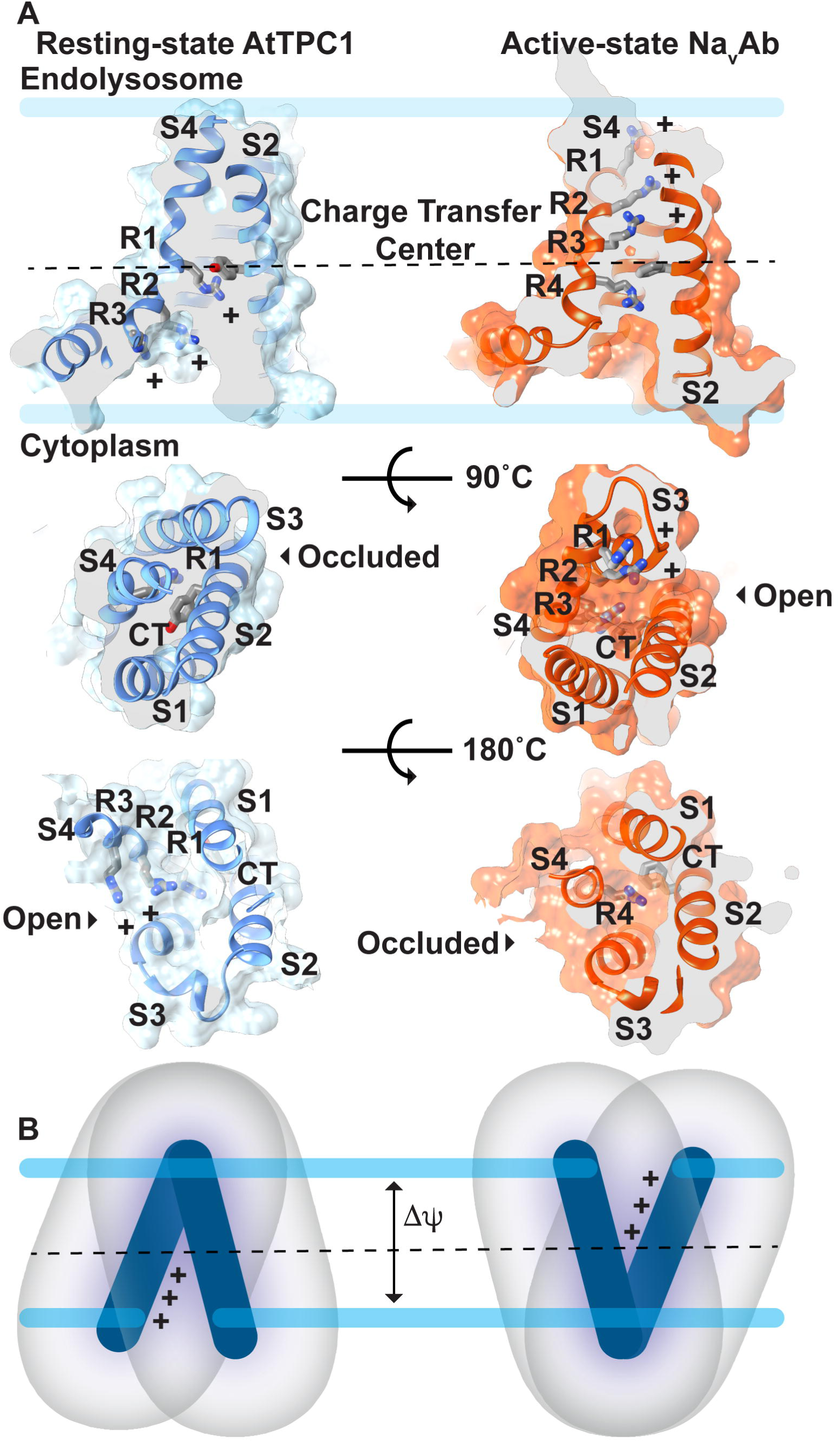
Figure 3. Mechanism of Voltage-sensing. A) Comparison of (left) resting-state AtTPC1 (PDB **5DQQ**; ref. [20]) and (right) active-state Na_v_Ab (PDB **3RVY**; ref. [56]) structures. View through the (Top) membrane, (Middle) luminal/extracellular, and (Bottom) cytoplasmic faces of the VSD. The charge transfer center (CT) location (dash line) in S2 and gating charges (R1-R4) in S4 are shown. B) A hypothetical pathway for gating charge movement in the VSD from the cytoplasmic leaflet in the resting-state through the membrane to the luminal leaflet in the active-state.

In AtTPC1, S10 is a 3_10_-helix. The fact that S10 maintains at least partial 3_10_-helicity in most active-state structures[56–59,62] and our resting-state suggests that the helical pitch does not change drastically during voltage-dependent activation. Rather 3_10_-helicity may be a conserved structural feature of voltage-sensors and required for function, as with K_v_ channels[57].

The M484L VSD2 mutation in human TPC2, located in S8 near the charge transfer center, is linked to melanomas and regulation of pigmentation[17,52]. The corresponding M479L mutation in AtTPC1 would disrupt the hydrogen bond with W505 in S9 and could perturb voltage-dependent activation. Structures of AtTPC1 in the active-state will shed light on the rearrangements required for channel activation.

Polyunsaturated fatty acids inhibit plant TPC1[50]. We observe a lipid bound to the interface between P1 and VSD2. This has been observed in previous structures of voltage-gated channels[63,56,64,65]. Van der Waals interactions between the lipid alkyl chain and M237 in S5 and L533 in S10 could stabilize the closed-resting state. A hydrogen bond between S277 in the S5-S6 pore loop and the fatty acid head group could also restrict pore opening. The electron density is consistent with palmitic acid, the most abundant fatty acid present in soy polar lipids–the major lipid source used for crystallization[20]. Lipid-binding to this site could restrict pore movement during voltage-dependent activation.

## Ion Selectivity

A primary function of most ion channels is to selectively conduct ions through the membrane in response to excitatory stimuli. This requires a mechanism to prevent incorrect ions from entering the channel lumen. In most cation-selective channels, a domain called the selectivity filter[66], formed by the pore loops, between S5 and S6, that line the channel, imparts selectivity through structural interactions with passing ions. Selective channels coordinate passing ions either through direct or water-mediated interactions[56,66,67]. The selectivity filter in AtTPC1 is asymmetric, with diameters across P1-P1’ and P2-P2’ of ~14Å and ~5Å, respectively. Residues with negative charge are present in the selectivity filter that could coordinate ions, but they are only aligned with the conduction pathway in P2-P2’. The negative charge density in the selectivity filter may explain the cation selectivity, but AtTPC1 may be unable to distinguish cations based on hydration shell as suggested for Na_v_s and Ca_v_s[56,67]. AtTPC1 is 5-fold selective for Ca^2+^-ions over other ions[68]. The structure identifies a binding site for Ca^2+^ replacement Yb^3+^-ion in the upper selectivity filter[20]. E605 directly chelates the Yb^3+^-ion, suggesting that this may provide some of the modest selectivity for Ca^2+^-ions. This site may also explain how trivalent lanthanides and Al^3+^-ions inhibit plant TPC1[69].

Mammalian TPC1/2 are Na^+^-selective (TPC1; P_Na_/P_K_~100, TPC2; P_Na_/P_K_~30) [40], although both can conduct Ca^2+^ under certain conditions[70]. Indeed exchange of the less selective AtTPC1 selectivity filter in P2 for human TPC2 produces a Na^+^-selective channel[68]. In the TPC2 selectivity filter, a second asparagine forms an additional constriction point that contacts passing Na^+^-ions[68]. Unlike other Na^+^/Ca^2+^-selective channels, which use negatively charged amino acid sidechains to interact with passing cations[56,67,71], TPCs use the uncharged sidechains of two asparagine residues. Therefore, TPCs use neutral amino acid sidechains for selecting Na^+^/Ca^2+^-ions—a distinct ion selectivity mechanism from other channels.

## Pharmacology

In animals, NED19 prevents infection by filoviruses either by altering fusion of viral membranes with the endolysosome, or by altering the binding of the viral glycoprotein to the NPC1 receptor in endosomes[19]. Serendipitously, NED19 was developed as an NAADP antagonist by virtual screening[72]. How NED19 in mammalian TPCs inhibits putative activation by NAADP is not clear. NED19 acts as a non-competitive inhibitor of NAADP activation by directly blocking mammalian TPC channels[73]. While AtTPC1 does not respond to NAADP, NED19 potently blocks channel conductance in endolysosomes (our unpublished data). The structure of AtTPC1 shows that the antagonist NED19 may prevent channel activation by clamping the pore domains to VSD2 (Figure 4). NED19 interacts with the luminal faces of P1 and P2. This suggests that NED19 may block channel opening by restricting movement of the pore helices P1 and P2.

**Figure 4.**
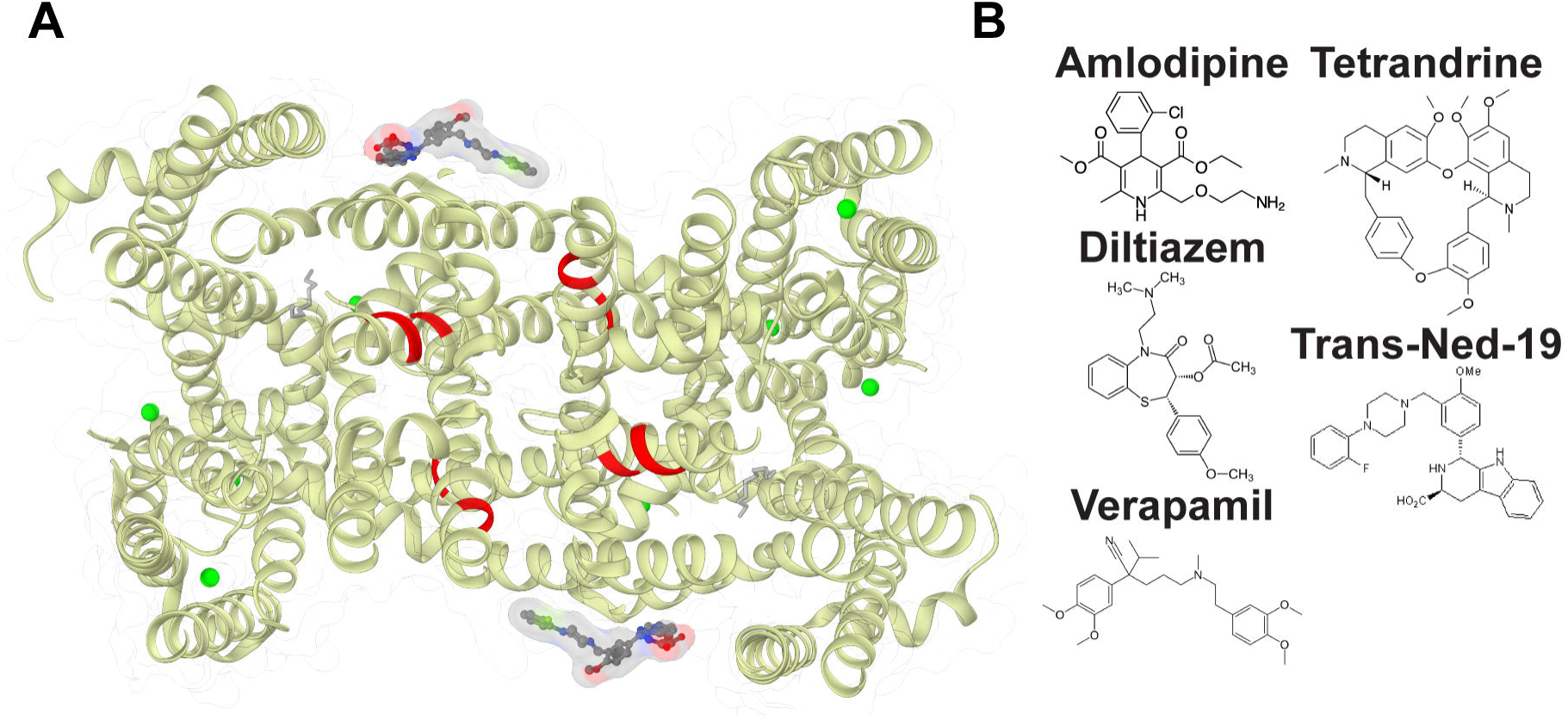
Figure 4. TPC Pharmacology. A) Predicted binding site for DHP (red) Ca_v_ blockers based on sequence homology with Ca_v_s overlaid onto the AtTPC1 structure bound to non-DHP molecule trans-NED-19 (PDB **5DQQ**; ref. [20]). B) Molecular structure of representative pharmacophores that block TPC channels.

Approved medications of the dihydropyridine (DHP), phenylalkylamine (PAA), and benzothiazepine (BTZ) classes of L-type Ca_v_ antagonists block Ebola virus trafficking and fusion in the endolysosome in a TPC1/2 dependent manner[19]. These drugs, like Tetrandrine and NED19, presumably act by blocking channel conductance. TPC channels contain conserved DHP and PPA binding sites in transmembrane segments S6 and S12 that may explain the pharmacological overlap with Ca_v_s[25,74,75]. Recent structures of bacterial Ca_v_ channels (Ca_v_Ab) bound to DHP (amlodipine and nimodipine) and PPA (verapamil) inhibitors suggest that inhibitors bind in two configurations[65]. DHP molecules bind to the pore periphery like NED19, whereas PPAs directly block the pore lumen. The pore asymmetry in TPCs may change the way these molecules, especially DHPs, link the pore and VSD domains.

Determining the antagonist binding sites on TPCs may serve as a template for the development of novel therapies for filoviral infections and possibly cancer.

Tetrandrine, a bis-benzylisoquinoline alkaloid isolated from the Chinese herb *Stephania tetrandra*[76], potently blocks TPC channels. NED19 and Tetrandrine both show promise in treating cancers[18] and filoviral infections[19]. Given the structural similarity to verapamil, an L-type Ca_v_ antagonist, Tetrandrine likely binds TPCs in a distinct location from NED19.

## Phosphoregulation by Kinases

Phosphoregulation is a hallmark of TPC channel function. Plant TPC1 is both activated (CAM-like dependent kinase) and inhibited (PKA, PKC, or a PKG-like kinase) at distinct sites by phosphorylation[77]. Cytoplasmic kinases modulate human TPC1/2 conductance and function[78–80], but only mTORC1 has been shown to directly inhibit channel currents through formation of high-affinity complexes[13].

We determined the location of phosphorylation sites in AtTPC1 in both the NTD (S22, T26, T29) and CTD (S701) (unpublished data; Figure 5). The NTD contacts EF1 via van der Waals interactions and hydrogen bonds. Since EF1 is not required for channel activation, the NTD phosphorylation sites may not directly gate the channel.

**Figure 5.**
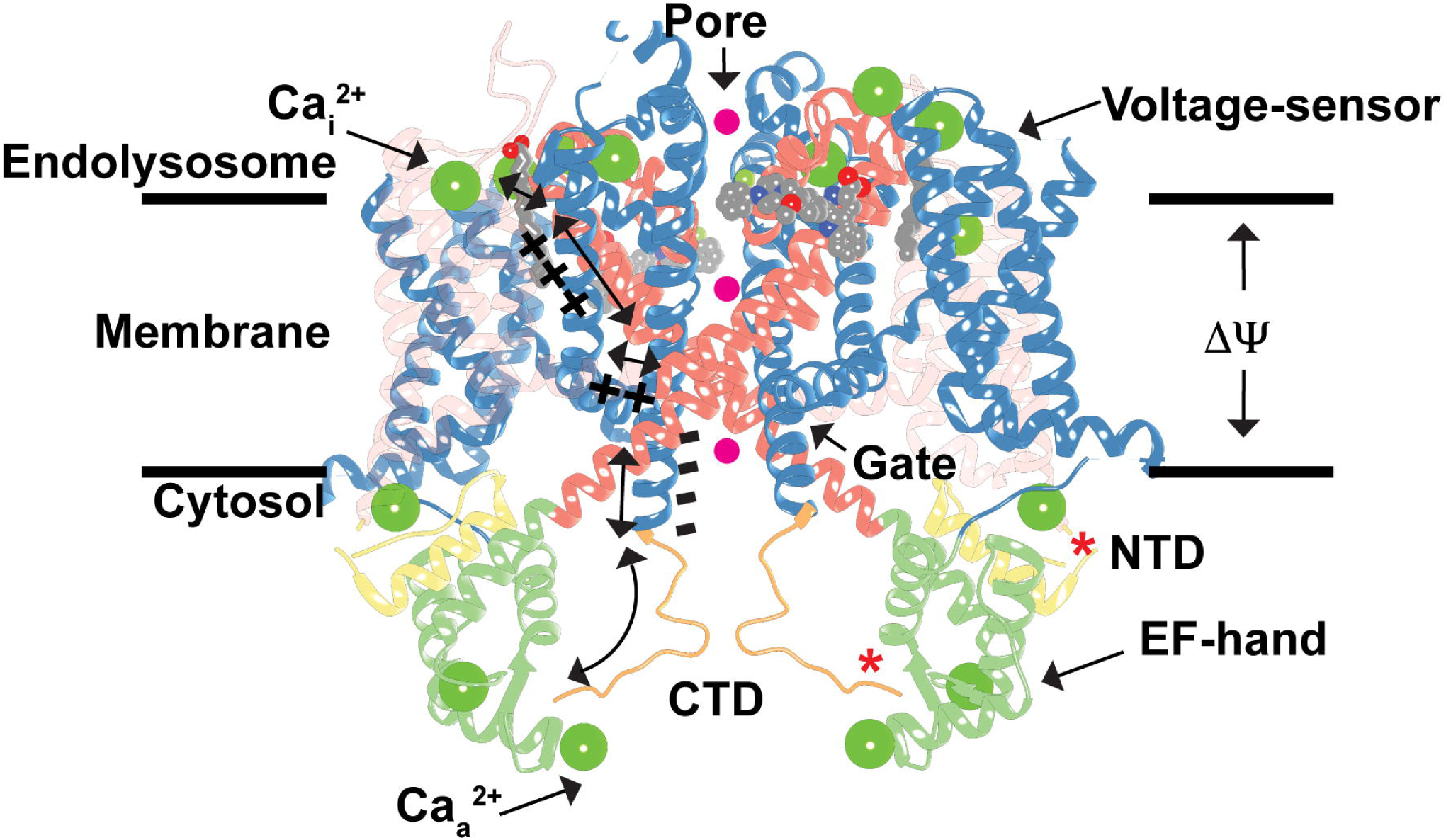
Figure 5. Model for TPC1 Activation and Regulation. Hypothetical model for TPC1 activation by membrane potential (**ΔΨ**) and cytosolic Ca^2+^ ions (**Ca_a_ ^2+^**), and inhibition by luminal Ca^2+^ ions (**Ca_i_^2+^**) and phosphorylation (**\***) overlaid onto the AtTPC1 structure (PDB **5DQQ**; ref. [20]). A possible mechanism for conformational coupling between voltage-sensor VSD2 and EF-hand domain EF2 during channel activation is shown. Hypothetical permeating ions are shown as spheres (magenta). The transparency of VSD1 is increased for clarity.

However, S701 in the CTD directly contacts the activating Ca^2+^-ion in EF2. Phosphorylation of S701 could disrupt the interaction of the CTD with EF2, preventing Ca^2+^-binding, thereby mimicking the apo-Ca^2+^ state. This may weaken the EF2-CTD interaction, increasing flexibility in the region, and thereby perturb the structural interaction between the CTD and the S10-S11 linker in VSD2. Pinpointing the sites of phosphorylation-dependent gating will be critical for defining how TPCs integrate kinase signaling to control the resting endolysosomal membrane potential and pH.

## TPC-protein Interactions

Human TPC1/2 form stable complexes with mTORC1 at the endolysosomal membrane that couple changes in cellular metabolism and nutrient concentrations to control of the endolysosome membrane potential and pH[13]. MTORC1 also forms complexes with Rag GTPases, Ragulator, SLC38a9 arginine transporter, NPC1 Niemann-Pick C1 cholesterol transporter, and H^+^-ATPase proton pumps[81–84] as part of the endolysosomal nutrient-sensing and transport machinery. These macromolecular complexes link dynamic cytoplasmic nutrient concentrations, cellular growth signals, and stress to changes in the endolysosome transport[85,86]. Uncovering the molecular interactions within this complex will open the way to understanding how mTOR signaling regulates endolysosome physiology.

The human TPC2 NTD contains a Rab GTPase-binding motif, not conserved in human TPC1. Rab-binding regulates trafficking and channel function[87]. The interaction of plant TPCs with Rab GTPases has not been tested, but the structure of AtTPC1 shows that binding to NTD could affect channel opening by influencing the structure of the nearby EF domains. While it is not known yet whether plant TPCs form protein-protein interactions via the NTD or CTD, the TPC1/2 CTD may form a hub for binding of apoptotic machinery[88].

We conclude that the structure AtTPC1 provides key insights into the mechanisms of voltage-sensing, ion selectivity, channel activation, and the phosphoregulation of TPC channels. It provides a template for the design of experiments probing the regulatory mechanisms of TPC channels.

## Acknowledgements

This work was supported by NIH grant GM24485 (to R.M.S) and a Postdoctoral Independent Research Grant (to A.F.K) from the University of California, San Francisco Program for Breakthrough Biomedical Research, which is partially funded by the Sandler Foundation. Beamline 8.3.1 at the Advanced Light Source is operated by the University of California Office of the President, Multicampus Research Programs and Initiatives grant MR-15-328599. The Berkeley Center for Structural Biology is supported in part by the National Institutes of Health, National Institute of General Medical Sciences, and the Howard Hughes Medical Institute. The Advanced Light Source is supported by the Director, Office of Science, Office of Basic Energy Sciences, of the US Department of Energy under Contract No. DE-AC02-05CH11231. Use of the Stanford Synchrotron Radiation Lightsource, SLAC National Accelerator Laboratory, is supported by the US Department of Energy, Office of Science, Office of Basic Energy Sciences under Contract No. DE-AC02-76SF00515. The SSRL Structural Molecular Biology Program is supported by the DOE Office of Biological and Environmental Research, and by the National Institutes of Health, National Institute of General Medical Sciences (including P41GM103393). Figures were made with Chimera developed by the Resource for Biocomputing, Visualization, and Informatics at the University of California, San Francisco (supported by NIGMS P41-GM103311).

